# Simulation and empirical evaluation of biologically-informed neural network performance

**DOI:** 10.1101/2025.11.13.687845

**Authors:** Gwen A. Miller, Ahmed Roman, Marc Glettig, Haitham A. Elmarakeby, Saud H. AlDubayan, Jihye Park, Ryan L. Collins, Eliezer M. Van Allen

## Abstract

Biologically-informed neural networks (BiNNs) offer interpretable deep learning models for biological data, but the dataset characteristics required for strong performance remain poorly understood. For instance, we previously developed P-NET, a BiNN with an architecture based on the Reactome pathway database, and applied this model to predict metastatic status of patients with prostate cancer using somatic mutation and copy number information. It seems likely that including additional relevant signal – e.g., germline variation in this context – should improve model performance, but we currently lack a principled approach to assess whether BiNNs will successfully detect this signal.

Here, we developed two simulation frameworks to evaluate the factors that influence BiNN performance – including signal type, signal strength, feature sparsity, and sample size – and empirically tested how integrating germline and somatic data affects the model’s ability to predict prostate cancer metastatic status. Simulations revealed that small sample size, weak signal strength, and especially extreme feature sparsity limit BiNN performance, and that the model preferentially uses linear over nonlinear signal. Empirically, P-NET performed poorly on sparse germline data, and while adding germline to somatic data did not improve prediction, it improved gene prioritization and model interpretation.

Broadly, our simulation frameworks enable systematic evaluation of how dataset-level characteristics affect BiNN performance and provide a principled framework for benchmarking novel methods.

## Introduction

New strategies for modeling multimodal molecular data that maximize both predictive performance and interpretability may advance biological discovery and biomarker development across cancer types. Among such strategies, biologically-informed neural networks (BiNNs) that merge principles of deep learning with known biology to enable interpretability have emerged as a widely employed approach^1–5^. These methods vary both in implementation and application, and potentially provide avenues to integrate seemingly disparate types of molecular information (e.g., multiple types of somatic features, somatic and germline combinations) and model states of interest (e.g., disease progression, treatment response).

For example, our group previously proposed P-NET, a BiNN built with a foundation of signaling data from Reactome^6^, and we examined its utility in classifying prostate cancers as primary or metastatic using a combination of somatic mutation and copy number data^7^. Through this approach, we demonstrated the potential of a model like P-NET both for biomarker development and for biological discovery. A subsequent reusability study validated P-NET’s usability and robustness, and systematically assessed the performance of several graph neural network architectures leveraging HumanBase tissue-specific gene-gene interaction data^8^. Other examples that incorporate prior knowledge into the model architecture include DeepOmix^5^, which incorporates signalling pathway databases and was used to predict patient survival from multi-omics data, and NeST-VNN^4^, which incorporates multiprotein assembly information and was used to predict drug resistance in breast cancer.

Our original application of P-NET was restricted to somatic data modalities. In prostate cancer, it has been previously established that germline features, such as pathogenic germline mutations in *BRCA2*, also contribute to metastatic progression^9^. Therefore, BiNNs that integrate both germline and somatic data are needed to maximize the potential of these approaches for precision oncology. Similarly, for P-NET and other BiNNs, investigators have begun exploring integration of additional molecular features, such as transcriptional or proteomic data^1,5,10,11^, even though we lack a fundamental understanding of the factors necessary for successful BiNN performance. Such factors include power for inference given cohort sizes and signal strength, as well as the stability of a BiNN’s interpretable outputs under various perturbations. Broadly, the ability to characterize the properties of a given molecular input type that contribute to strong BiNN performance could serve as a valuable guide for the responsible and effective use of these models in research and perhaps inform future clinical deployment.

Here, we address this knowledge gap regarding the use of BiNNs for precision oncology generally, and the relative merits of combining somatic and germline data for prostate cancer risk classification (primary vs. metastatic) using P-NET in particular, using a combination of simulations and empirical assessments. Our simulations reveal that small sample size, weak signal strength, and especially extreme feature sparsity categorically limit BiNN performance. Deeper interrogation of these simulations further suggest that BiNNs preferentially use linear over nonlinear signal. For prostate cancer risk classification, we find that P-NET performs poorly on the germline data alone – which is extremely sparse – and that adding germline to somatic data does not improve prediction. That said, adding germline data does improve gene prioritization in the model’s interpretable outputs; this shows that germline data contains additional information relevant to metastatic disease prediction, even if it is too sparse to directly improve predictive performance *per se*.

## Results

### Development and exploration of a simulation framework for biologically-informed neural network model performance

We developed a framework for simulating the performance of a BiNN (e.g., P-NET) for a classification task between two sets of biological samples, labeled X_0_ and X_1_. (Fig. 1A). We first created three types of signal, assigning genes a perturbed mean (linear signal), membership in class-specific correlated modules (nonlinear signal), or both.

**Figure 1.**
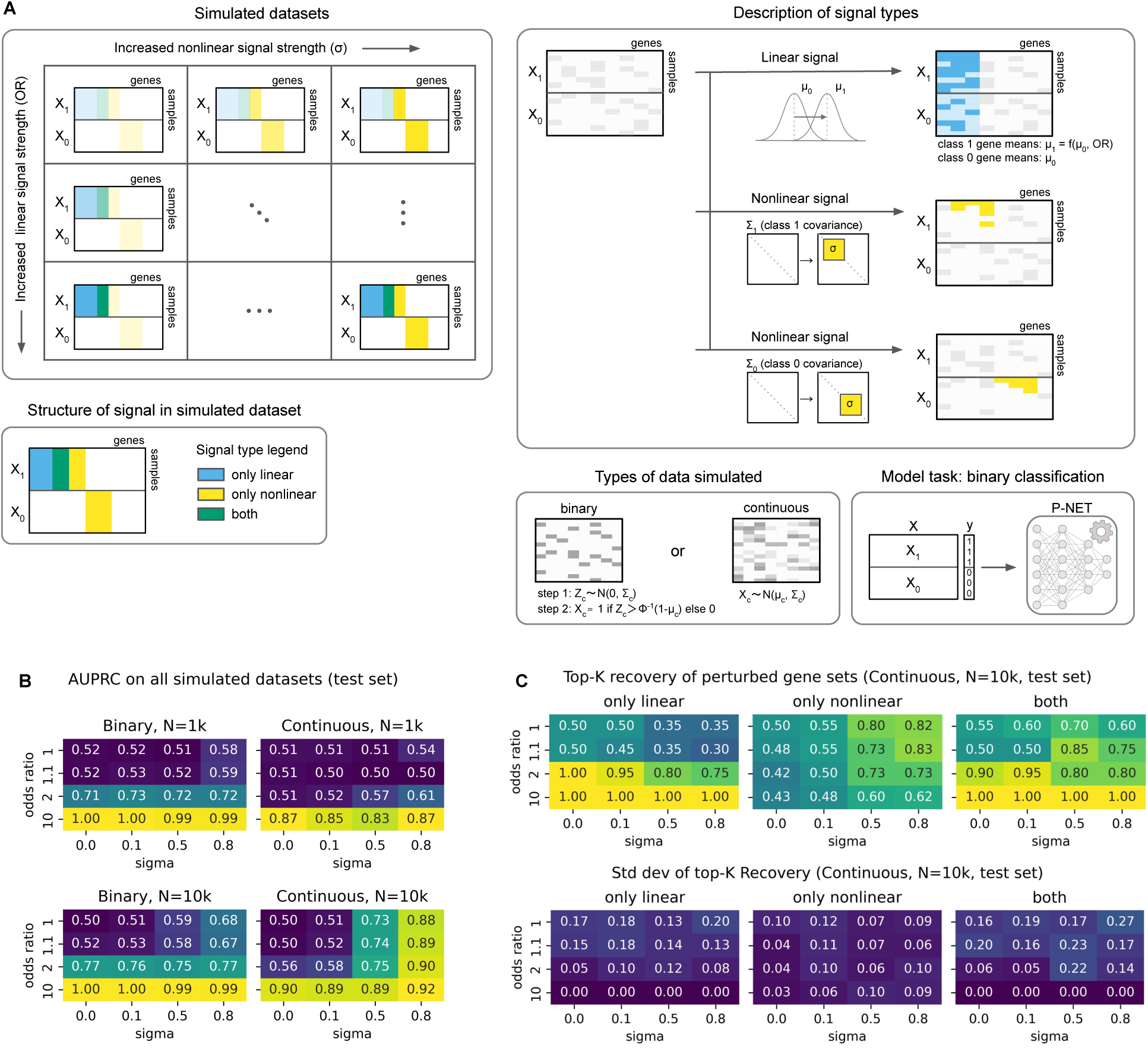
P-NET performance on simulated datasets with varied signal type and strength. **A** We generate simulated datasets along two axes of perturbation: linear signal strength and nonlinear signal strength (“Simulated datasets”). We inject linear signal into a subset of genes by assigning a larger mean in class 1 compared to class 0 according to the specified odds ratio (OR), which determines the strength of the linear signal. Nonlinear signal is injected with class-specific gene-gene correlation modules. Genes in a module share correlation with strength sigma (σ); the rest are assigned near-zero correlation using Gaussian noise (“Description of signal types”). Each simulated dataset contains some genes with only linear signal, only nonlinear signal, or both as illustrated in “Structure of signal in simulated dataset”. Data are sampled either as continuous (multivariate normal) or binary (thresholded latent Gaussian) values using class-specific sampling parameters: mean vectors (μ₀, μ₁) and covariance matrices (Σ₀, Σ₁) (“Types of data simulated”). P-NET’s binary classification performance is assessed on each simulated dataset. **B** Performance (test AUPRC) across simulated datasets with varied odds ratio (linear signal strength) and σ (nonlinear signal strength). Results are shown for binary and continuous data with N=1k and N=10k samples. Values reflect the mean over n=10 replicates. **C** Top-K recovery (continuous data, N=10k) of three types of perturbed gene sets: genes with only linear signal, only nonlinear signal, or both. The first row of heatmaps contains the mean top-K recovery while the second row contains the standard deviation over n=10 replicates.

Linear signal strength was defined by the odds ratio (OR) of each gene in X_1_ versus X_0_. Nonlinear signal strength was defined by correlation strength (σ). We systematically explored how signal type and strength impact model performance by varying the OR and σ. This framework also supports generation of both continuous and binary datasets, e.g., where a gene’s mean could correspond to average expression or mutation frequency, respectively.

We then evaluated the area under the precision-recall curve (AUPRC) of P-NET across varying simulation regimes (Fig. 1B). Model performance improved as linear signal strength (OR) increased, and P-NET’s ability to detect nonlinear signals was especially dependent on sample size. With N=1k samples, increasing nonlinear signal (σ) did not improve performance when linear signal was held constant, except for continuous data at OR=2. With a 10-fold increase in simulated sample size (N=10k), increasing σ generally improved AUPRC, except when the model already performed well at σ = 0. For continuous data, even with equal class means for every gene (OR=1), increasing σ increased the AUPRC from 0.50 (σ=0) to 0.88 (σ=0.8). This demonstrated that, as expected, BiNNs are capable of relying solely on nonlinear signal.

We next utilized this simulation framework to understand how different types of signal are used by BiNNs. We computed top-K recovery of gene sets where either the mean, the correlation structure, or both differed between classes (Fig. 1C, Fig. S1). We observed that recovery of genes with linear signal increased with large OR or small σ. Increasing σ improved recovery of genes with nonlinear signal at N = 10k, though impact was diminished when performance was already strong at σ = 0, and had no effect at N = 1k (Fig. 1C, Fig. S1A–C), mirroring our earlier findings. Without nonlinear signal (σ=0), increasing gene linear signal between classes led to improved recovery of genes. Without linear signal (OR=1), increased σ improved the recovery of genes with nonlinear signal. Of note, recovery of genes with only nonlinear signal decreased as OR increased. For example, P-NET could detect nonlinear signal at σ=0.5 in continuous data with 10k samples, yet used less nonlinear signal when performance is dominated by linear signal; recovery of genes with only nonlinear signal dropped from 0.8 (OR=1) to 0.6 (OR=10).

Thus, the simulation framework demonstrated that BiNNs did not always use signals they were powered to detect. Finally, BiNN performance strongly correlated with improved model stability, as evidenced by decreased standard deviation in predictive performance and gene recovery (Fig. 1C, Fig. S1). Taken together, this framework showed that BiNNs can leverage both linear and nonlinear biological signals, exhibit a bias toward linear signal utilization, and yield interpretable outputs aligned with their predictive behavior.

Next, to dissect the interaction between linear signal strength and feature sparsity with more representative background properties, we developed a second simulation framework based on observed somatic mutation data from a previous prostate cancer genomics dataset (hereafter “P1000” from Armenia, et al. 2018^12^). We randomly split the samples into a control class (X_0_) and a perturbed class (X_1_), then applied single-gene perturbations to X_1_, as summarized in Figure 2A. We controlled linear signal strength with OR, and feature sparsity with the control frequency, which was defined as a gene’s mutation rate in class 0.

**Figure 2.**
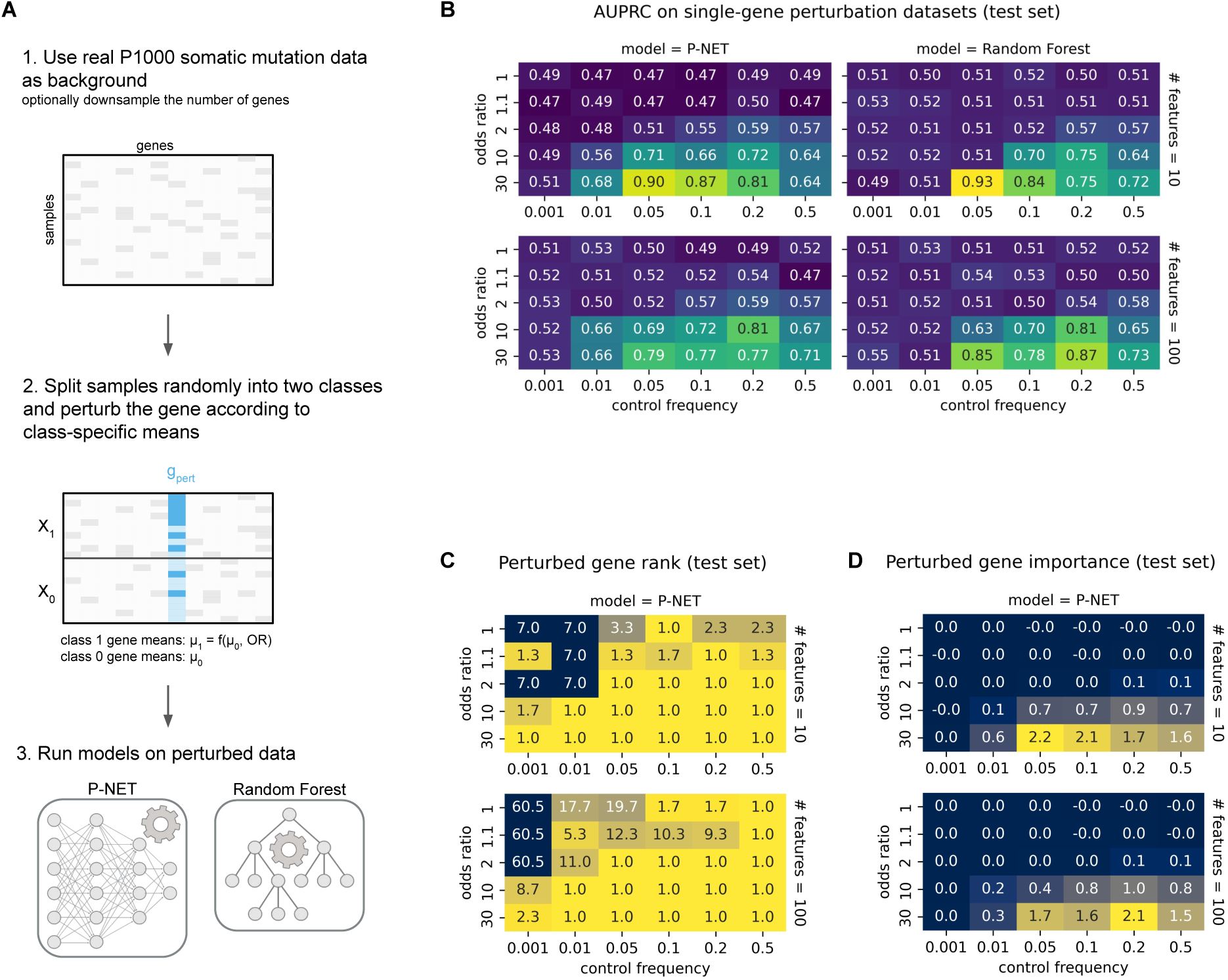
Single-gene perturbation on observed P1000 somatic mutation data. **A** Simulation framework. We select one gene to perturb by increasing its mutation probability in class 1 according to a target odds ratio. All other genes are unmodified. Control frequency (μ₀) defines the feature sparsity and odds ratio defines the perturbation strength. We perform binary classification on the perturbed data using P-NET and random forest (RF). **B** Test set AUPRC on all tested combinations of odds ratio and control frequency. Columns show results for P-NET and RF; rows correspond to different numbers of retained features (# features = 10 or 100). **C** P-NET’s ranking of the perturbed gene across all sampling conditions. **D** P-NET’s assigned feature importance for the perturbed gene across the same conditions. All values in **B-D** reflect the mean over n=3 replicates.

Performance of P-NET and random forest (RF) was similar across a range of simulation parameters (Fig. 2B). Both models showed increased AUPRC with increased ORs for linear signal, and best performance with control frequencies between 0.05 to 0.2. Generally, larger control frequency was needed to detect weaker signals, as expected due to decreased statistical power for rare perturbations. P-NET outperformed RF when control frequency was low, and extreme sparsity prevented detection even of strong signals in both models (for example, when control frequency = 0.001). Overall, BiNNs like P-NET appear to have a lower threshold for detecting true signal than simpler models like RF, as shown by the boundary separating random from non-random performance across datasets varying in signal strength and feature sparsity.

Increasing the number of retained non-informative features (# features = 100 vs 10 genes) did not significantly affect test set performance (Fig. 2B). However, there was slight overfitting in runs where 100 non-informative features were retained and/or the data were modeled with RF, as evidenced by nonrandom performance on the train set in the absence of ground-truth signal (OR = 1, AUPRC near 0.6) (Fig. S2). Only P-NET showed the expected random performance on the train set in the absence of ground-truth signal (# features = 10, Fig. S2).

Finally, we evaluated whether P-NET correctly prioritized the perturbed gene in two ways: average rank of the gene (Fig. 2C) and its assigned gene importance score (Fig. 2D), where lower rank or larger importance scores both indicated better BiNN signal detection. In all models with non-random performance, the perturbed gene was ranked first (Fig. 2B, 2C). Gene rank was not useful when model performance was near random, as gene importance values were near zero: apparent prioritization did not reflect true signal. For example, in simulations with no true signal (OR = 1), prioritization of the perturbed gene increased with control frequency (Fig. 2C). Thus, BiNN performance strongly correlated with perturbed gene importance and rank.

### Empirical assessment of BiNN performance on multimodal data

The simulation frameworks gave us a principled way to assess the impact of several key factors on BiNN behavior, including signal detection thresholds, predictive performance, and interpretable outputs. However, simulations are inherently limited in their ability to capture the complexity and nonrandom error modes inherent to real-world, multi-modal molecular datasets.

Thus, to complement simulation-based examination of BiNNs, we empirically examined how the inclusion of multiple molecular data modalities affected BiNN performance in a clinically relevant setting. Given the prior use of the P1000 somatic cohort with P-NET for prediction and discovery of genes involved in prostate cancer metastasis^7^, and the known importance of germline genetic features in prostate cancer development and progression^9,13^, we chose to evaluate P-NET’s ability to predict metastatic status using paired somatic and germline data from the P1000 dataset (Fig. 3A). We trained P-NET and RF models with somatic data only, germline data only, or a combination. The somatic data included three modalities: mutations, amplifications, and deletions. We constructed various germline datasets from combinations of pathogenic rare, common, missense, and loss-of-function variants, as summarized in Figure S3.

**Figure 3.**
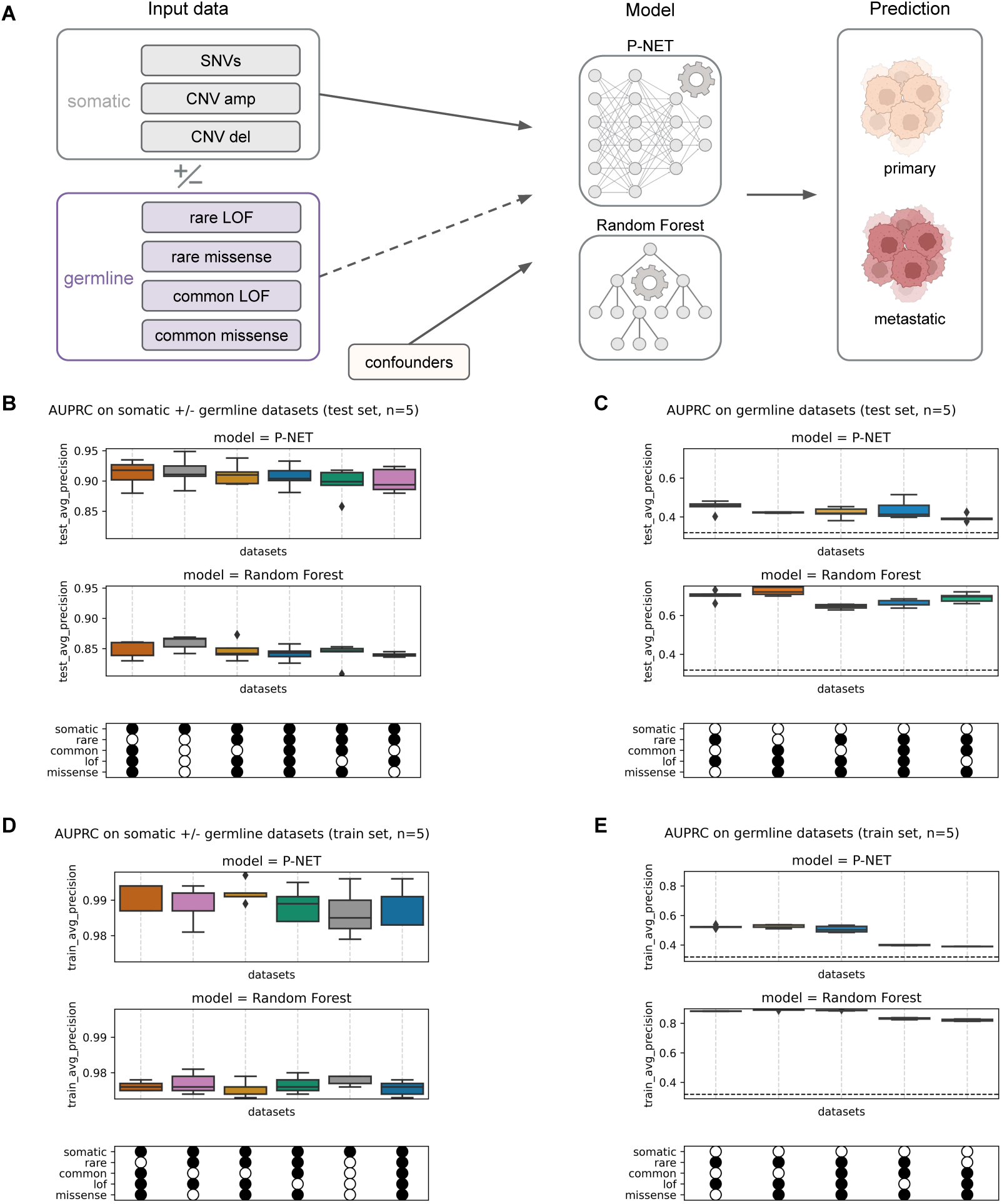
Performance on varied combinations of P1000 somatic and germline datasets. **A** Schematic of the empirical task setup, where we evaluate models on their ability to predict metastatic status using paired somatic and germline data from the P1000 dataset. We train models with somatic data only, germline data only, or a combination. We construct various germline datasets from combinations of pathogenic rare, common, missense, and loss-of-function variants. AUPRC of P-NET vs. random forest (RF) on **B** somatic and varied germline data, evaluated on the test set and **C** varied germline data, evaluated on the test set. **D** and **E** are equivalent to **B** and **C** except evaluated on the train set. Random performance is represented with a black dashed horizontal line in **C** and **E** at the prevalence of the positive (metastatic) class. The x-axis legend encodes which datasets were passed to the model, with a filled circle meaning inclusion. The first row indicates whether the 3 somatic modalities were included (mutation, amplification, and deletion). The next four rows describe which germline variants were included: rare, common, LOF, or missense variants. All boxplots in **B-E** show the median AUPRC +/- one quartile over n=5 replicates.

P-NET performance was similar on somatic and combined somatic + germline datasets, and better than RF (average AUPRC ≥0.89 and ≤0.86 respectively; Fig. 3B). Adding germline data did not improve performance of either model.

On all combinations of just germline data, RF outperformed P-NET (Fig. 3C). P-NET’s low performance on the train set (AUPRC≤0.5) suggests that this model underfit the data compared to RF (train AUPRC≥0.8) (Fig. 3E). RF exhibited greater overfitting than P-NET, reflected in a larger train–test AUPRC gap (Fig. 3B vs. 3D; Fig. 3C vs. 3E). P-NET’s low performance is likely due to data sparsity; samples contain on average less than one pathogenic germline mutation (Fig. S3C).

Lastly, we examined how inclusion of an additional stream of molecular data affected the model’s interpretable outputs. As expected based on prior research^9^, the addition of germline data substantially improved the importance ranking of *BRCA2* for primary vs. metastatic classification. While *BRCA2* was ranked in the 8th percentile (388 of 4854 genes) when using somatic data alone, its rank improved to the 1.6th percentile (78/4854) in the combined somatic + germline setting (Fig. 4). And though P-NET performed poorly on germline data alone, *BRCA2* still ranked first overall in gene importance based on germline data that included rare loss-of-function variants (Fig. 4). Thus, addition of complementary molecular signals that capture different genomic information can modify the importance assigned to biologically relevant genes, even if overall BiNN predictive performance remains constant. To facilitate comparison to the simulation results in Figure 2, additional tables with gene rank and empirically-calculated OR and control frequencies are available in Figure S4.

**Figure 4.**
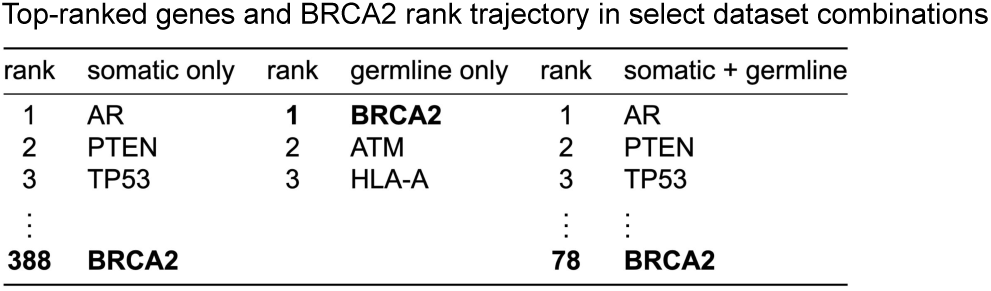
Interpreting P-NET results for select combinations of P1000 somatic and germline datasets. Rank trajectory of BRCA2 under three data conditions: somatic only, germline only (rare LOF), and combined somatic + germline. Top three genes shown for each condition; BRCA2’s rank is highlighted. The reported gene rank was derived by taking P-NET-derived gene importances, calculating per-run gene rank, averaging these ranks across n=5 replicates, and rank-ordering the result.

## Discussion

Overall, this study served two primary goals: simulation and empirical assessment of biologically-informed neural network (BiNN) performance. First, we developed two complementary simulation frameworks to systematically evaluate the performance of BiNNs. The first let us assess the impact of signal type, signal strength, and sample size. The second let us assess the interaction between linear signal strength, feature sparsity, and the number of noninformative features.

Using P-NET as our representative BiNN, we found that signal detection was inhibited by small sample size, weak signal strength, and feature sparsity. Extreme sparsity prevented detection even of strong signals, a limitation observed in both BiNNs and traditional models like RF. Detecting nonlinear signal required larger sample size and was inhibited when performance was dominated by linear signal. This point is particularly relevant because analyses integrating multiple data modalities are of interest; however, cross-dataset signals are inherently nonlinear (e.g., germline and somatic double-hit events observed for many canonical tumor suppressor genes^14–17^) and therefore especially difficult for BiNNs to detect in settings with limited data.

Moreover, the three categories of perturbed genes allowed us to benchmark models not only by predictive performance (e.g., AUPRC), but also by their ability to recover and attribute signal to the underlying perturbation. P-NET’s top-ranked genes corresponded to those with signal, and performance correlated strongly with gene importance scores. This suggests that P-NET’s performance came from correctly prioritizing relevant features.

The single-gene spike-in simulation results provide a lookup table for linear signal detection thresholds as a function of signal strength (OR) and prevalence (control frequency) when using BiNNs (e.g., Fig. S4). Users can calculate odds ratios and control frequencies for features in their dataset and compare them to the detection thresholds derived from these simulations. If no features exhibit detectable linear signal, strong nonlinear effects or larger sample size may be required for non-random model performance. In addition, the simulations can be adapted to new models and new datasets with minimal code changes. Running the simulations on new models enables direct benchmarking against baselines such as P-NET and RF. The single-gene perturbation simulations can also be rerun using alternative observed datasets as background. This enables more accurate explanations for model behavior by aligning the simulation environment more closely with the characteristics of the dataset under study.

To complement this simulation framework and compare with real-world performance, we also evaluated P-NET’s ability to predict metastatic status using paired somatic and germline data from the P1000 dataset. We found that P-NET performed poorly – and significantly worse than RF – on extremely sparse data, such as on germline data alone. Under less sparse data conditions, the performance of P-NET eclipsed RF. Adding germline data to somatic data did not improve predictive performance, but shifted gene rankings, indicating that BiNNs may provide improved interpretations even in the absence of classification gains.

Empirical and simulation signal detection thresholds were similar with respect to linear signal strength and feature sparsity. P-NET’s top-ranked genes on empirical data were characterized by OR and control frequency values that correspond to nonrandom performance in single-gene spike-in simulations. P-NET detected sparser signals than RF in our spike-in simulations.

However, we saw the opposite when we tested the models on germline data, where RF outperformed P-NET. One potential explanation for this contradiction lies in the simulation design. These spike-in simulations generated a single sparse gene using the somatic mutation dataset as background, whereas every gene in the germline dataset is extremely sparse.

Rerunning the spike-in simulations using germline mutation data as background could resolve whether dataset-specific characteristics are responsible for this apparent discrepancy.

While our simulation framework offers useful guidance, several limitations should be considered when generalizing these results to new datasets, tasks, and models. The simulation results will overestimate performance on smaller datasets and underestimate performance on larger datasets than we tested here. We only assessed binary classification tasks; BiNN behavior in regression or multiclass settings remains to be explored. We observed consistent trends in signal detection and overall performance in P-NET and RF, suggesting that the factors we assessed may have similar impact in other models.

Broadly, this work introduces a simulation framework for examining the factors underlying BiNN performance. One key finding from both simulations and empirical tests was that model performance was poor on sparse data. Yet we know that sparse data can contain relevant signals; e.g., prior work showed that germline mutations are enriched in metastatic vs. primary prostate cancer^9^. Therefore, a fruitful direction for future work would be the development of novel methods capable of performing well on sparse data.

Expanding the size of existing datasets may aid performance of existing methods, especially in datasets with sparse features or primarily nonlinear signal. Developing methods capable of integrating multiple datasets – e.g., germline data from multiple cancer types – could also help overcome small sample sizes. Combinations of these innovations may further molecular discovery and biomarker development across a range of real-world clinical scenarios.

## Methods

### Simulating datasets using a joint distribution

### Defining perturbed gene sets

We first simulated genetic data by sampling from class-specific joint distributions designed to reflect structured mean and correlation patterns across genes (see Fig. 1A). For each class (class 1 and class 0), we defined a vector of per-gene means μ_c_ (denoted μ_1_ and μ_0_) and a corresponding gene-gene covariance matrix (Σ_1_ and Σ_0_). Baseline gene means μ₀ were each set to 0.1 as this matched some of the highest frequencies we saw in the binarized P1000 datasets, though the simulation code enables sampling uniformly from a predefined range. To introduce signal, a subset of genes were assigned an odds ratio (OR) > 1 to simulate association with class 1; their alternative class frequencies μ₁ were calculated from μ₀ using the definition of OR. For remaining genes, we set μ_1_=μ_0_ (equivalent to OR=1).

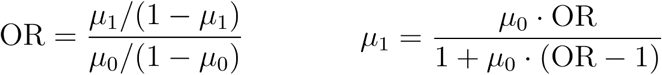

To model gene–gene dependencies, we embedded a “module” of genes with elevated pairwise correlation σ into each class-specific correlation matrix. Values for genes not in the correlation module were assigned according to a Gaussian N(0, 0.01^2^) to represent background noise. The final covariance matrices were derived by scaling the correlation matrices by user-provided gene standard deviations (default 1).

### Defining gene sets with ground-truth signal

To simulate realistic patterns of association, we designed the simulation to produce three non-overlapping gene sets with distinct types of ground-truth signal. First, we perturb a set of genes so their mean values differ between class 0 and class 1, representing marginal differences in feature distribution. Next, we constructed class-specific correlation modules by selecting two disjoint sets of genes (one for each class) to form tightly correlated blocks within the class-specific covariance matrices. Importantly, we ensured that half of the genes in the class 1 correlation module were also drawn from the Δμ set.

This design yields three interpretable categories of signal:

1. linear signal: genes where only the mean differs between classes (Δμ-only),
2. nonlinear signal: genes where only the correlation structure differs (correlation-only), and
3. both linear and nonlinear signal: genes where both the mean and correlation structure differ (both Δμ and correlation).

This modular structure allows us to benchmark models not only by predictive performance, but also by their ability to recover and attribute signals to the correct type of perturbation.

### Generating the datasets from class-specific distributions

We used these two parameters, μ_c_ and Σ_c_, to generate continuous or binary data on a per-class basis. To generate continuous data (e.g., log-transformed gene expression), we sampled from a multivariate normal distribution N(μ_c_, Σ_c_). To simulate binary genotype data (e.g., mutation status), we employed a latent Gaussian thresholding approach. First, latent variables were sampled from a multivariate normal distribution with zero mean and a class-specific covariance matrix encoding the desired gene–gene correlations. Then, each latent value was thresholded at the standard normal quantile corresponding to 1-μ_c_, ensuring that the probability of assignment to 1 matched the specified marginal probability μ_c_; values above the threshold were set to 1, and values below were set to 0.

Specifically, we generated datasets containing 100 genes. Of these, 20 had a perturbed mean (μ_1_ ≠ μ_0_). The Σ_1_ module contained 20 genes, 10 of which also had a perturbed mean. The Σ_0_ module contained 20 non-overlapping genes. Each simulated dataset had the same number of samples per class. We tested all combinations of: number of samples per class ∈ {500, 5000}, OR ∈ {1, 1.1, 2, 10}, σ ∈ {0, 0.1, 0.5, 0.8}, and data type ∈ {binary, continuous}. The simulation process allowed us to systematically vary signal strength (via OR) and the correlation structure (via σ) to evaluate the robustness and interpretability of predictive models.

### Simulating datasets with single-gene spike-in perturbation

To benchmark model performance as a function of linear signal strength (OR) and feature sparsity (control frequency), we generated simulated binary genotype datasets with single-gene perturbations (see Fig. 2A). Starting from an observed binary genotype matrix (P1000 somatic mutations) to maintain realistic gene-gene correlation structure and background noise, we selected a single target gene (AR) for perturbation. The perturbed gene’s mutation frequency in class 1 (μ_1_) was calculated from the mutation rate in class 0 (i.e., control frequency; μ₀) using the definition of OR.

For each simulation, we split samples randomly into two classes of equal size. For each class, the specified proportion of samples (μ_c_) was randomly assigned 1 (mutation) in the target gene column; all other samples were assigned 0 (no mutation). The rest of the genotype matrix was left unchanged, except for optional random subsampling of non-target genes to control feature dimensionality. When downsampling, we ensured that the perturbed gene was included in the final matrix. The rest were chosen at random. We tested all combinations of: OR ∈ {1, 1.1, 2, 10, 30}, control frequency ∈ {0.001, 0.01, 0.05, 0.1, 0.2, 0.5}, and number of retained features ∈ {10, 100}.

### Preparing somatic and germline data

#### P1000 somatic and germline data

We used tumor and matched germline prostate cancer patient whole-exome sequencing (WES) samples assembled by Armenia et al.^12^ We used the same somatic mutation and copy number data as in the P-NET paper^7^, which were derived from a unified computational pipeline for harmonized somatic alteration derivation (n=1,013 patients)^12^. Initial processing of the germline data (n=1,072 patients) was performed in AlDubayan et al. and included stringent quality control, detection of germline variants with DeepVariant^18^ (v0.6.0), functional annotation with Ensembl’s Variant Effect Predictor (VEP, v92.0), and genetic ancestry inference via principal component analysis^19^. We restricted our analyses to the 943 patients with paired somatic and germline WES data, and will refer to this as the “P1000 cohort”.

### Defining a germline gene subset

We restricted the analysis of germline data to variants within a predetermined gene set (n=824 genes) containing three groups of genes implicated in cancer risk: somatic (n=575), germline tier 1 (n=143), and germline tier 2 (n=106). The somatic genes were selected from the Catalog of Somatic Mutations in Cancer (COSMIC) Cancer Gene Census^20^ (CGC, https://cancer.sanger.ac.uk/census, v86).

The germline cancer risk genes were compiled from COSMIC CGC germline genes and selected publications (Rahman et al., Huang et al., Mirabello et al.)^20–23^. Genes with sufficient supporting evidence based on manual evaluation by S.H.A. were assigned to germline tier 1; others were assigned to tier 2. Specifically, all CGC germline tier 1 genes were automatically assigned tier 1 here. For the remaining genes, the number of published case-control papers showing an association between the gene and any cancer determined the gene’s status: genes that showed sufficient gene-cancer association in two or more studies were assigned to tier 1, genes shown to be associated with a cancer phenotype in only 1 study were assigned to tier 2, and genes without published gene-cancer phenotype evidence were excluded.

### Filtering of pathogenic germline variants

The process for filtering the germline VCF to just pathogenic variants is illustrated in Supplementary Figure 3. First, we filtered out any remaining low-quality variants using quality thresholds based on minimum depth (DP≥10), minimum Phred score (GQ≥20), and minimum variant allele frequency (VAF≥0.25). Then, we processed these variants based on their clinical significance annotation in the Clinical Variation database (ClinVar) (https://www.ncbi.nlm.nih.gov/clinvar/, June 2021)^24^ and Genome Aggregation Database population frequencies (gnomAD)^25^. We removed variants annotated as Benign/Likely Benign (B/LB) and retained variants annotated as Pathogenic/Likely Pathogenic (P/LP). We also retained variants with conflicting ClinVar annotations if there were at least as many P/LP submissions as existed in any of the other categories (B, LB, Uncertain). We retained variants that lack a ClinVar annotation if they had a severe consequence (i.e., a stop codon, a frameshift, a canonical splice site variant, or other similar variants defined as VEP impact = HIGH) and minor allele frequency (MAF) <0.01 in gnomAD. We also removed variants that occurred in more than 5% of our dataset’s samples as these are likely technical artifacts. The result was a pathogenic germline samples-by-variants matrix.

### Pathogenic germline variant matrix to various gene-level datasets

We considered subsets of these pathogenic germline variants along two axes: variant prevalence and variant impact. Variants with MAF<0.01 were considered “rare”, and the rest “common.” We considered variants with high impact and moderate impact as predicted by VEP, which we refer to as LOF (loss-of-function) and missense throughout the manuscript as these variant types constitute the bulk of each impact category.

Next, we generated input datasets with various combinations of prevalence (rare vs. common) and impact (LOF vs. missense). First, we selected all variants that match the desired characteristics (e.g., rare LOF and rare missense variants). Second, since P-NET expects inputs at the gene level, we collapsed variants into genes and binarized (e.g., if a sample has any variant in BRCA2, record 1, otherwise 0). The result was a binary samples-by-gene matrix which can be passed into machine learning models. We generated the following combinations to aid assessment of which types of pathogenic germline variants contain the most predictive signal:

1. rare LOF (equivalent to all LOF in our setting because no common LOF variants exist in the P1000 pathogenic germline data)
2. all missense (rare missense + common missense)
3. all rare (rare LOF + rare missense)
4. all common (equivalent to common missense since no common LOF variants exist in the P1000 pathogenic germline data)
5. all (rare LOF + rare missense + common missense)

### Harmonizing somatic and germline data inputs

Our data harmonization script loaded the somatic and germline mutation datasets, confounders, and prediction target. We restricted the data to samples with both somatic and germline data. As in Elmarakeby et al., the somatic data included all genes expressed in TCGA prostate or genes in the Reactome pathway database. The germline datasets contain only a subset of these genes, as described earlier. Since P-NET restricts all data modalities to overlapping input features, we chose to zero-impute the missing genes to enable the retention of all somatic data used in the original P-NET paper.

We ran P-NET and RF on various combinations of the P1000 data: somatic only, germline only, or a combination of both. The top 10 genetic ancestry principal components were always included as confounders to account for the impact of population structure.

### Model training and evaluation

#### Model hyperparameters

For both simulations and the empirical analysis, we used P-NET’s default settings^7^, which were originally optimized for the P1000 somatic dataset. The only modification was increasing the number of training epochs from 400 to 500 (with early stopping enabled) to allow sufficient time for convergence.

When we ran RF on the empirical task, we set min_samples_split = 50 (around 5% of samples) and used default values for all other parameters. To reduce RF overfitting in the single-gene spike-in simulations, we selected hyperparameters that minimized the gap between training AUPRC and validation AUPRC. Using Weights and Biases^26^, we performed a random search over max_depth ∈ {2, 3, 5, 7, 10, 15} and min_samples_leaf ∈ {1, 5, 10, 50, 100} on datasets with control_frequency = 0.1, n_features = 10, and odds_ratio ∈ {1, 2, 10, 30}. Across all odds ratios, min_samples_leaf had a stronger impact than max_depth, with the optimal combination being max_depth = 2 and min_samples_leaf = 100.

### Data splits

For the joint simulations, we used a random 70%/15%/15% train/validation/test split. For the empirical task, we applied the same split used in the original P-NET study: an 80%/10%/10% train/validation/test split of 1,013 patients. Restricting to the 943 patients with matched germline data, this corresponded to an 80.5%/9.4%/10.1% split (759/89/95 samples). We applied the same split for the single-gene spike-in simulations, which used the P1000 somatic mutation data from these same 943 patients as a background.

### Training replicates

To account for variability due to model stochasticity, we repeated each model run multiple times using different random seeds. We used 10 seeds for the joint simulation experiment, 3 seeds for the single-gene spike-in simulation experiment, and 5 seeds for the empirical task.

### Model evaluation

The prediction performance was measured using AUPRC, AUC, F1, accuracy, and balanced accuracy. We report the mean and standard deviation across replicates for all data splits (train, validation, and test): joint simulation (n=10, Tables S1-2), single-gene spike-in simulation (n=3, Tables S3-4), and empirical (n=5, Tables S5-6). In the main text, we focus on the AUPRC.

For the empirical task, we also report the OR and control frequency for the top 10 ranked genes in a subset of tested data inputs: somatic only, all germline (rare LOF + rare missense + common missense), and somatic + all germline (Fig. S4).

## Data and Code Availability

Custom code was developed as part of the analysis reported here, and has been deposited on GitHub: https://github.com/vanallenlab/pnet-simulation.

The somatic data was sourced from the original P-NET paper^7^, and can be located here: https://zenodo.org/records/10775529. The germline raw sequence data (BAM files) can be obtained through dbGaP (https://www.ncbi.nlm.nih.gov/gap) as described in Supplementary Table 1 from Armenia et al.^12^. The germline data used in this manuscript can be generated based on Armenia et al.^12^ and AlDubayan, S.H. et al.^19^ with our methods section and code.

## Acknowledgements

This work was supported by NIH P50CA272390 (E.M.V.), NIH 1P01CA228696 (E.M.V.), DOD Prostate Cancer Data Science Award W81XWH-21-PCRP-DSA (E.M.V.), DOD CDMRP Award HT9425-23-1-0023 (H.A.E.), and the Mark Foundation Emerging Leader Award (E.M.V.). Some figures contain icons created in BioRender^27^ (Figs. 1A, 2A, 3A). During the preparation of this work the authors used ChatGPT in order to edit the manuscript for clarity and readability. After using this tool, the authors reviewed and edited the content as needed and take full responsibility for the content of the published article.

## Disclosures

Eliezer Van Allen:

Advisory/Consulting: Enara Bio, Manifold Bio, Monte Rosa, Novartis Institute for Biomedical Research, Serinus Bio, TracerBio

Research support: Novartis, BMS, Sanofi, NextPoint

Equity: Tango Therapeutics, Genome Medical, Genomic Life, Enara Bio, Manifold Bio, Microsoft, Monte Rosa, Riva Therapeutics, Serinus Bio, Syapse, TracerDx

Travel reimbursement: None Speaking Fees: TD Cowen

Patents: Institutional patents filed on chromatin mutations and immunotherapy response, and methods for clinical interpretation; intermittent legal consulting on patents for Foaley & Hoag Editorial Boards: *Science Advances*

---

**Figure S1.**
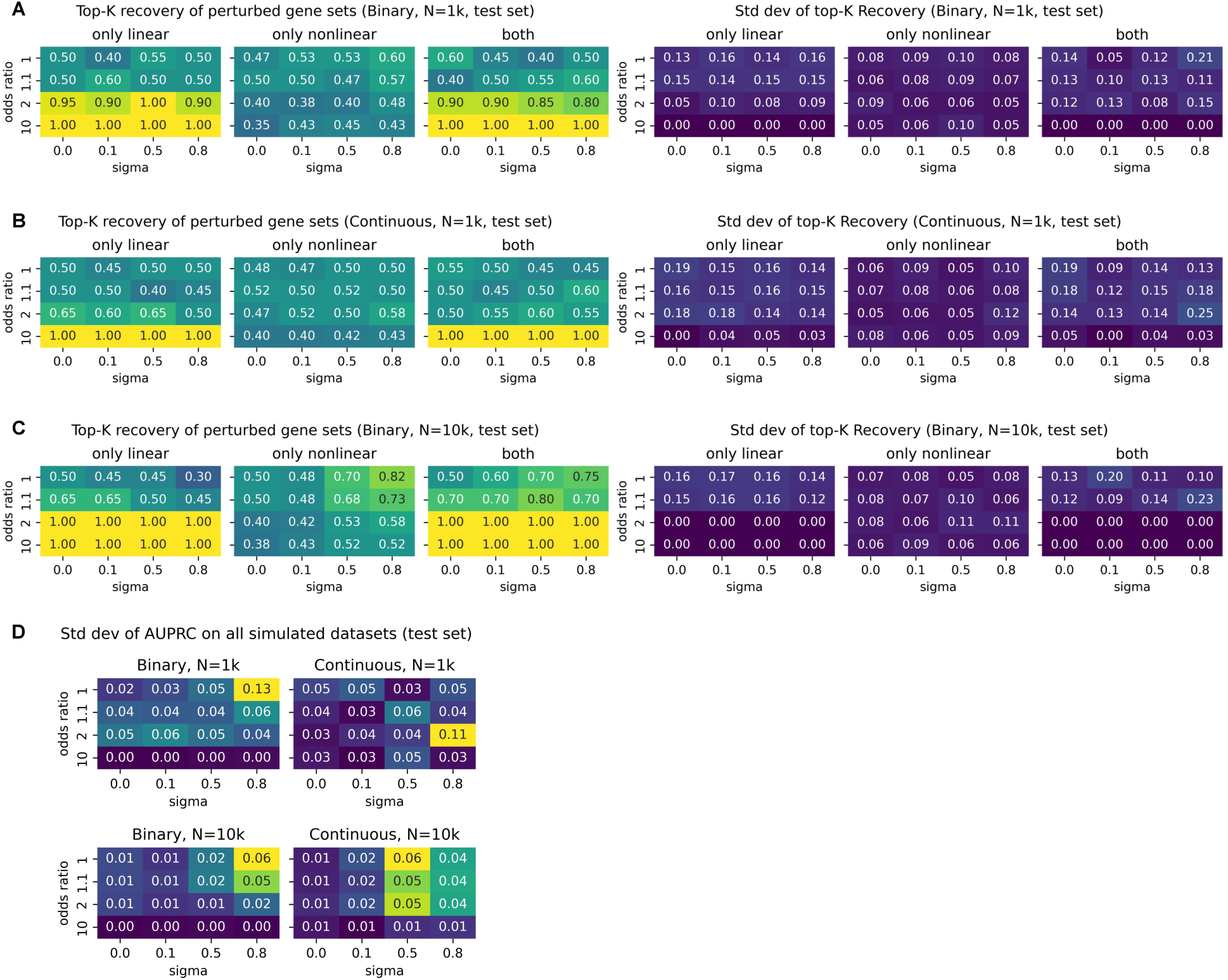
Top-K recovery of P-NET in simulated datasets with varied signal type and strength. Each subfigure shows the top-K recovery of three ground-truth gene sets across all tested combinations of odds ratio and sigma: genes where only their mean differs between classes (only linear), genes where only the correlation structure differs (only nonlinear), and genes where both differ. The title of each subfigure indicates the sampling strategy (binary or continuous) and the sample size (1k or 10k): **A** Binary sampling, N=1k **B** Continuous sampling, N=1k **C** Binary sampling, N=10k. The left three columns of heatmaps in **A-C** contain the mean top-K recovery while the right three columns contain the standard deviation over n=10 replicates. **D** Standard deviation in test AUPRC (n=10 replicates) across simulated datasets with varied odds ratio (linear signal strength) and σ (nonlinear signal strength). Results are shown for binary and continuous data with N=1k and N=10k samples.

**Figure S2.**
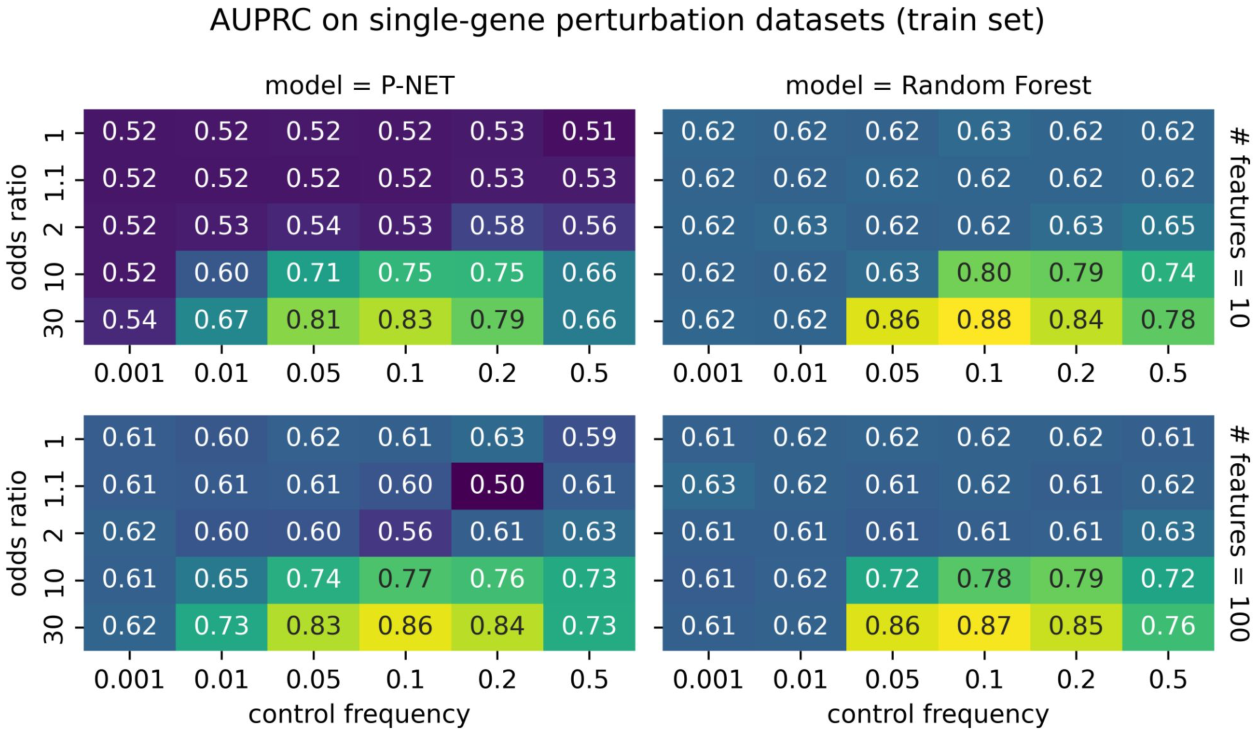
Single-gene perturbation on P1000 somatic mutation backbone evaluated on train set. Heatmaps report the train set AUPRC for each pair of sampling parameters, odds ratio and control frequency. In order, the subpanel columns report results from P-NET and random forest models. The subpanel rows correspond to the number of retained gene features (10 vs. 100, respectively). All values reflect the mean over n=3 replicates.

**Figure S3.**
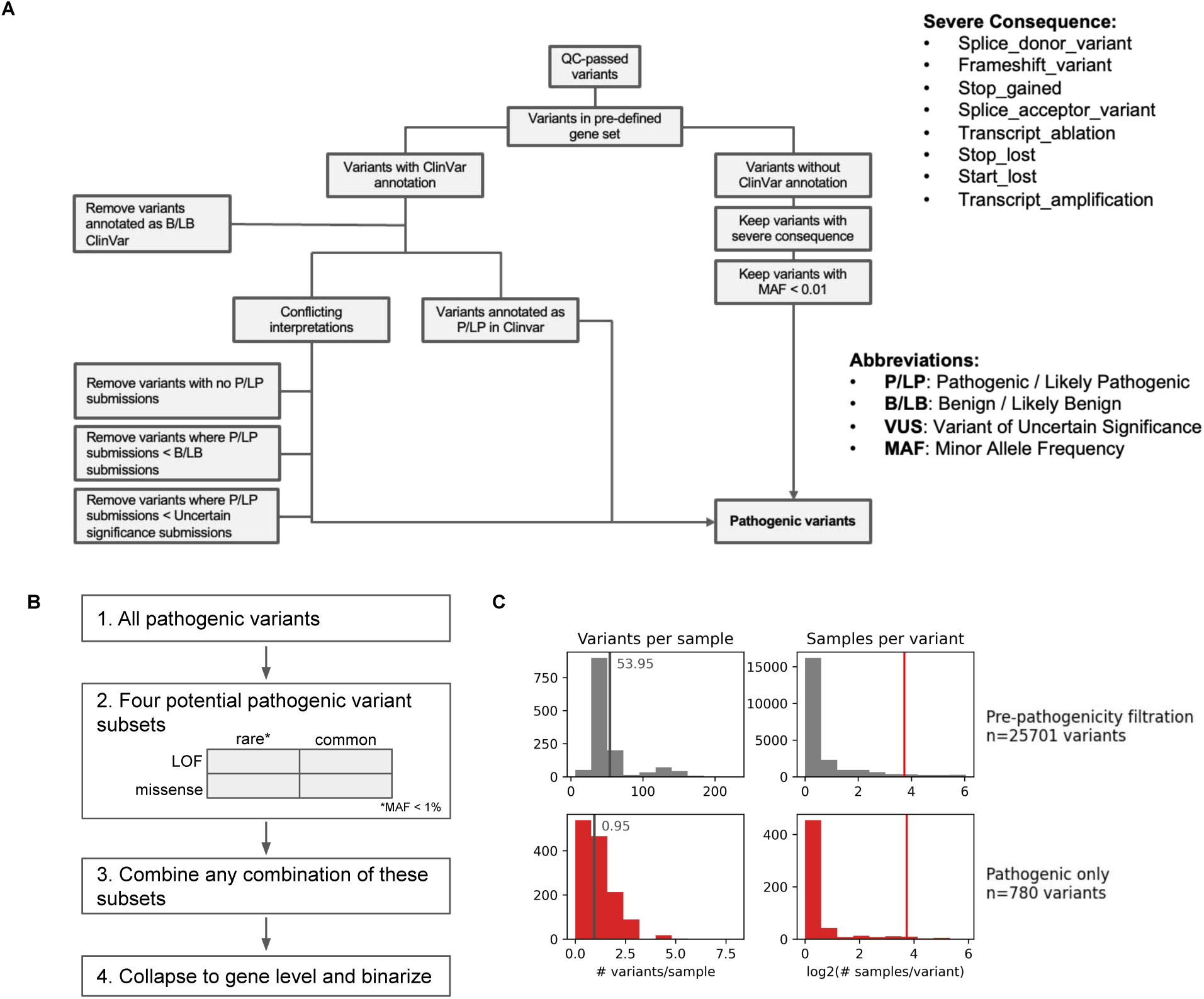
Details on curation of germline datasets. **A** Flowchart describing process for filtering a VEP-annotated germline VCF to just pathogenic variants. **B** Process for generating various gene-level pathogenic germline datasets (e.g., restricted to rare LOF mutations, or common LOF + common missense mutations). Rare is defined as a minor allele frequency (MAF) < 0.01 in gnomAD. **C** Histograms of variants per sample and samples per variant before and after pathogenicity filtering (first and second columns, respectively). The vertical grey line marks the average number of variants per sample. The vertical red line marks the number of samples corresponding to a dataset-level MAF=0.01.

**Figure S4.**
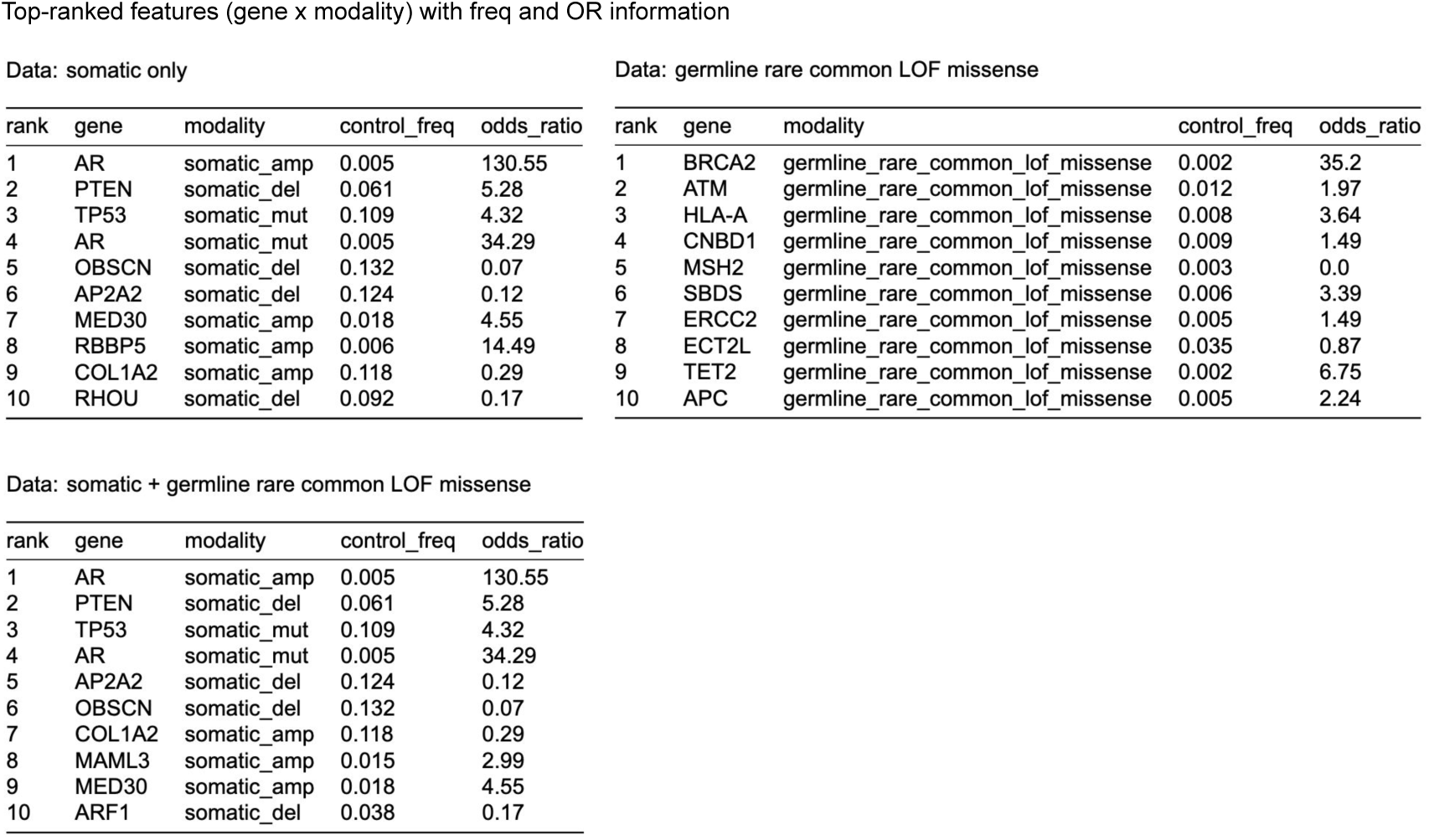
P-NET’s top-ranked features (gene x modality) with the corresponding odds ratio (OR) and control frequencies as calculated from the input datasets. Results reported for three different sets of input data: somatic only, all pathogenic germline LOF and missense variants (rare and common), and their combination.

